# *Bacillus subtilis* strain UD1022 as a biocontrol agent against *Magnaporthe oryzae,* the rice blast pathogen

**DOI:** 10.1101/2025.03.13.642990

**Authors:** Timothy Johnson, Lainey Kemmerer, Nalleli Garcia, Jessie Fernandez

**Author notes:** Equal contribution (Timothy Johnson, Lainey Kemmerer).

## Abstract

Rice blast disease, caused by *Magnaporthe oryzae*, is a major threat to global rice production, necessitating sustainable disease management strategies. Compared to chemical pesticides, biocontrol agents, such as beneficial microbe antagonists, offer a sustainable approach to naturally inhibit plant pathogens. This study evaluates the biocontrol potential of *Bacillus subtilis* UD1022 against *M. oryzae* through both direct antagonism and volatile-mediated inhibition. In dual culture assays, UD1022 significantly inhibited fungal growth. Furthermore, a stacking plate assay demonstrated that UD1022 produces volatile organic compounds (VOCs) that suppress fungal growth. Beyond vegetative growth inhibition, UD1022 also disrupted key infection processes of *M. oryzae*. Spore germination was reduced by 50%, while appressorium formation decreased by approximately 44% in UD1022-treated samples. *In planta* assays revealed that UD1022-treated rice plants exhibited a substantial reduction in disease severity compared to untreated controls. This reduction correlated with the upregulation of key defense genes in the salicylic acid, jasmonic acid, and ethylene signaling pathways, suggesting that UD1022 primes systemic resistance in rice plants. These findings establish UD1022 as a potent biocontrol agent capable of suppressing *M. oryzae* through direct antagonism, VOC-mediated inhibition, and induction of systemic resistance. This study underscores the potential of UD1022 as an eco-friendly alternative to chemical fungicides for managing rice blast disease.

**IMPORTANCE:** *Magnaporthe oryzae* is a destructive fungal pathogen that causes rice blast disease, leading to significant yield losses and threatening global food security. Here, we investigated the biocontrol potential of *Bacillus subtilis* UD1022, a beneficial rhizobacterium known for its plant growth-promoting and antifungal properties. Our *in vitro* and *in planta* studies revealed that UD1022 suppresses *M. oryzae* through direct antagonism, volatile organic compound (VOC)-mediated inhibition, and the induction of systemic resistance in rice. These findings demonstrate UD1022 as a promising candidate for microbial-based disease management and the role of beneficial bacteria in enhancing crop protection. This research contributes to the development of sustainable agricultural practices by leveraging naturally occurring microbes to improve plant resilience and disease resistance.

## INTRODUCTION

Rice (*Oryza sativa*) is one of the most vital crops for human nutrition, serving as a primary food source for over half of the world’s population. As population size increases and global demand for rice rises, sustainable and efficient agricultural practices are essential to ensure stable production. However, plant diseases pose a major challenge to rice cultivation, significantly threatening yields and food security (Nalley et al., 2016). Among the most destructive is rice blast disease, caused by the fungal pathogen *Magnaporthe oryzae* (syn. *Pyricularia oryzae*). This pathogen infects all above-ground parts of the rice plant, leading to severe yield losses (1, 2). Responsible for over 30% of annual harvested rice losses, rice blast is one of the most persistent and devastating threats to global food security, causing an estimated $66 billion in crop losses each year—enough to feed 60 million people (3, 4). Under favorable conditions—such as high humidity, frequent rainfall, and optimal temperatures ranging from 25–28°C for infection and lesion development—*M. oryzae* spreads rapidly. When these conditions persist, the pathogen can cause severe outbreaks, sometimes leading to total crop failure with yield losses reaching 100% (5).

*M. oryzae* infection begins when a conidium attaches to the leaf’s surface and germinates, producing a germ tube that forms an appressorium at its tip (1, 6, 7). The appressorium is a specialized pressure-generating structure that enables the fungus to penetrate host tissue by mechanically rupturing the tough leaf cuticle. It achieves this by generating enormous turgor pressure, which is directed through a narrow penetration structure at its base (8). Appressorial adhesion to the host cell is a crucial step in this process, facilitated by mucilage production, which enhances attachment and ensures successful infection. Once inside, *M. oryzae* develops invasive hyphae (IH), which invaginate the plant membrane to form the extra-invasive hyphal membrane (EIHM) compartment, a defining feature of its biotrophic phase that separates fungal structures from the host cytoplasm (9, 10). Another key component of this phase is the biotrophic interfacial complex (BIC), a specialized structure positioned initially at the tip of the primary IH and later relocating subapically in bulbous IH (11, 12). The BIC is essential for the targeted secretion of effectors, which are small fungal proteins that manipulate plant immune responses to promote successful colonization. As the infection progresses, the pathogen transitions to a necrotrophic phase, leading to the formation of necrotic lesions (13). Within these lesions, *M. oryzae* undergoes sporulation, producing new conidia that can be dispersed by wind or rain, allowing the disease cycle to continue and spread to new host plants. Managing rice blast disease is challenging due to *M. oryzae*’s genomic plasticity and wide geographic distribution, allowing it to rapidly evolve and circumvent host resistance through transposable elements and extrachromosomal circular DNAs (14–16). Its adaptability and hemibiotrophic lifestyle undermine current control strategies, rendering resistant cultivars ineffective and reducing fungicide efficacy, ultimately posing environmental and health risks (17). To sustain rice production, alternative and more sustainable disease management strategies must be integrated with existing approaches for long-term control.

In recent decades, biological control strategies have emerged as a promising alternative to synthetic pesticides and fungicides. Biological control agents (BCAs) are living organisms or their derived components that suppress pest or pathogen populations, typically through antagonistic interactions or host priming mechanisms (18, 19). Against fungal pathogens, BCAs—often bacteria or fungi—produce secondary metabolites, antibiotics, or other bioactive compounds with fungistatic or fungicidal properties (18). Depending on the target pathogen and crop system, BCAs can be applied as soil amendments, foliar sprays, or seed coatings (20).

Beyond direct antagonism, many bacterial BCAs promote induced systemic resistance (ISR) in host plants, enhancing plant immunity through cross-kingdom signaling and priming a more robust defense response (19, 21). Some BCAs also promote plant growth independently of pathogen presence, functioning as plant growth-promoting rhizobacteria (PGPR) by improving nutrient availability, root architecture, or plant hormone production (18). Among these, many *Bacillus* species, including certain strains of *Bacillus subtilis*, exhibit strong potential as both BCAs and PGPRs (22).

One such strain, *Bacillus subtilis* UD1022 (hereafter referred to as UD1022), originally obtained from Dr. Harsh Bais at the University of Delaware, has been commercialized for its ability to enhance plant growth and provide disease protection across various crop species (23). UD1022 is a Gram-positive, exopolysaccharide (EPS)-producing soil bacterium with a fully sequenced genome (24, 25). It produces small-molecule antimicrobial compounds, including the cyclo-peptide surfactin, and forms robust biofilms that contribute to its biocontrol function (23).

In addition to its PGPR benefits, UD1022 has demonstrated antagonistic activity against several phytopathogens, including *Colletotrichum trifolii, Ascochyta medicaginicola, Phytophthora medicaginis*, and *Clarireedia jacksonii* (25, 26). However, its potential to control rice pathogens, particularly *M. oryzae*, remains unexplored. Given its effectiveness against other economically significant fungal pathogens, this study investigates UD1022’s antagonistic interactions with *M. oryzae* and the mechanisms underlying its inhibitory effects. Through both *in vitro* and *in planta* assays, we demonstrate that UD1022 exhibits strong antagonistic activity against *M. oryzae*, likely mediated by a combination of direct antifungal mechanisms—including the production of volatile and non-volatile compounds—and ISR-induced plant defenses. These findings showcase UD1022’s potential as an effective biocontrol agent in the rice blast pathosystem and its role in integrated disease management strategies for sustainable rice production.

## RESULTS

### In vitro inhibition of M. oryzae by B. subtilis UD1022

To evaluate the biocontrol potential of UD1022 against *M. oryzae*, we conducted dual culture and volatile-mediated inhibition assays using the wild-type fungal isolate, Guy11. In a dual culture assay, UD1022 significantly inhibited fungal growth by 60%, whereas the negative control *Escherichia coli* DH5α showed no inhibition (**Fig. 1A-C**). Similarly, untreated fungal cultures exhibited unrestricted growth, confirming the specificity of UD1022’s antifungal activity. These results establish UD1022 as a highly effective antagonist against *M. oryzae*.

**Fig 1.**
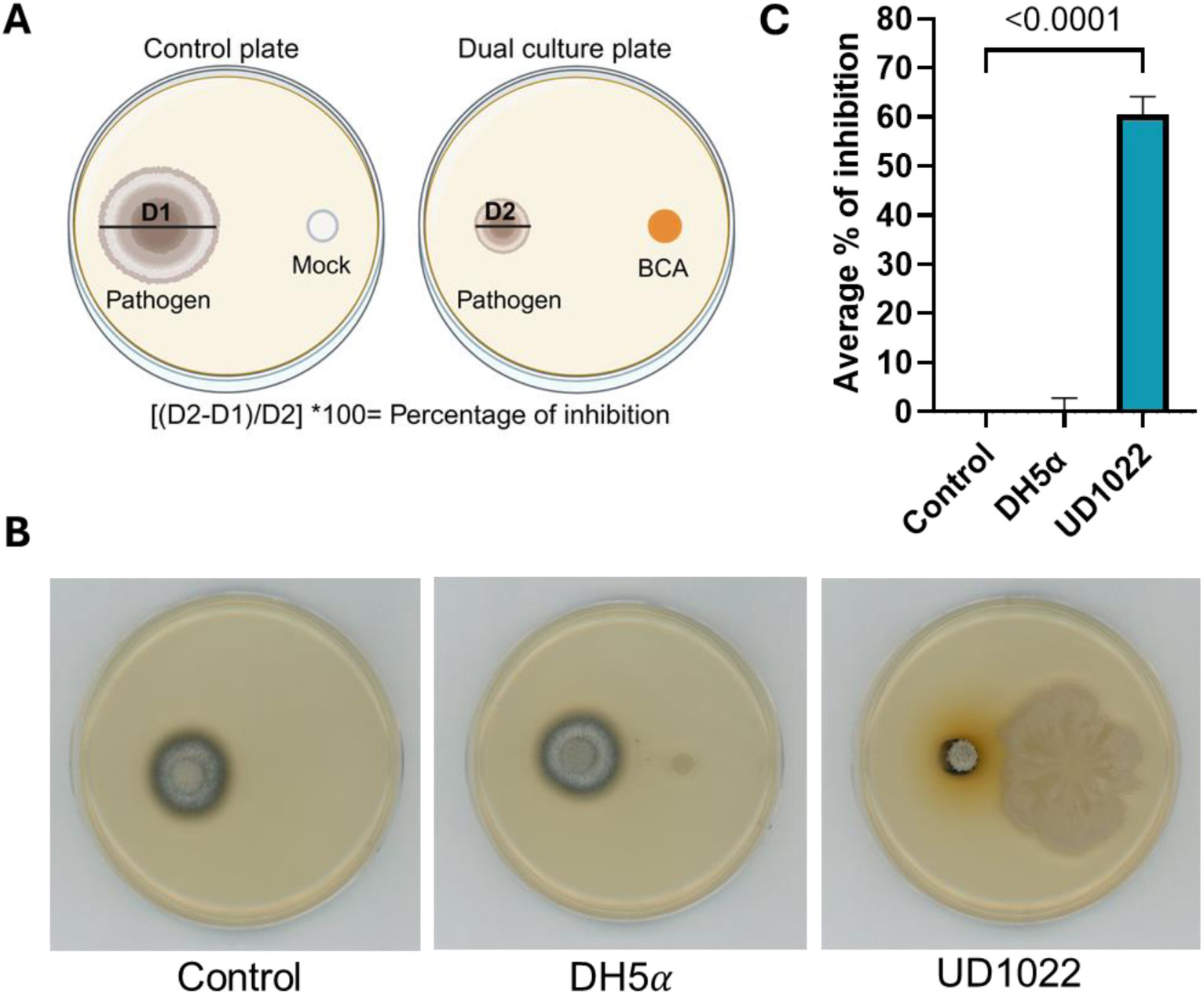
Antagonistic activity of *B. subtilis* UD1022 against *M. oryzae*. **A.** Dual culture assay CM plates. **B.** Diagram of the dual culture assay, where the control plate contains the pathogen with a water-control, and the dual culture plate contains the pathogen with the biocontrol agent. D1/D2 represent the pathogen’s diameter **C.** Percentage of fungal growth inhibition in the presence of UD1022 and control samples (DH5α and water-control). Plates were incubated at 28°C for 5 days. Images and measurements were taken at 5 days post incubation (dpi).

Once UD1022 was confirmed to exhibit direct antagonism, its ability to secrete volatile organic compounds (VOCs) was evaluated using the stacking plate approach (**Fig. 2A**). VOCs, often secondary metabolites, are known to suppress fungal pathogens, although previous studies reported that UD1022 did not inhibit *C. jacksonii* growth through VOCs generation (25). In contrast to these findings, our results demonstrate that UD1022 generates VOCs capable of inhibiting *M. oryzae* growth by approximately 30%, while the negative control *E. coli* DH5α and untreated control showed no *M. oryzae* growth inhibition (**Fig. 2B,C**). The combined results of antagonistic and volatile assays highlight UD1022 as a promising candidate for managing *M. oryzae* in rice crops, capable of suppressing fungal growth through both direct inhibition and VOC-mediated mechanisms.

**Fig 2.**
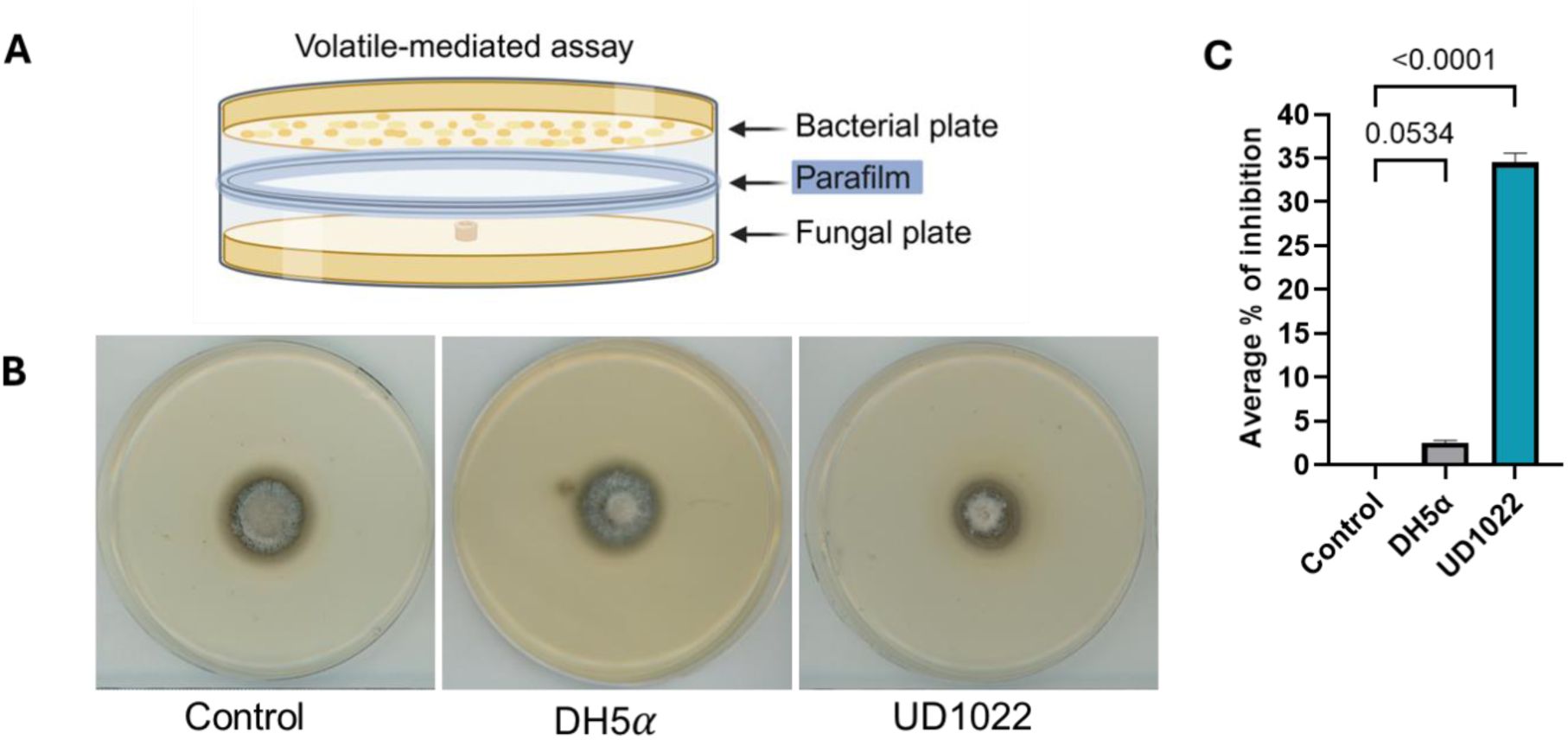
Volatility assay of *B. subtilis* UD1022 against *M. oryzae*. **A.** Stacking assay using CM plates. **B.** Diagram illustrating the volatile-mediated assay using a stacked plate method. **C.** Quantification of the percentage of fungal growth inhibition in the presence of UD1022 and control samples (DH5α and water-control). Plates were incubated at 25°C for 5 days. Images and measurements were taken at 5 dpi.

### Inhibition of spore germination and appressorium formation

To evaluate the inhibitory effects of UD1022 on the germination and appressorium formation of *M. oryzae* spores, we conducted assays using hydrophobic glass coverslips (**Fig. 3**). Coverslips were used to simulate the hardness and hydrophobicity of plant surfaces, promoting *M. oryzae* spore germination and appressorium development. Bacterial suspensions were combined with fungal spores, placed on glass coverslips, and observed for spore germination and appressorium formation. Results showed that *M. oryzae* spore germination was inhibited by UD1022, with an approximate 50% reduction compared to control treatments (*E. coli* DH5α and untreated spores) (**Fig. 3A**). Additionally, appressorium formation was reduced by approximately 44% at 20 hours post-inoculation (hpi) on coverslips treated with *B. subtilis* at different concentrations (**Fig. 3B; Fig. S1**). These findings demonstrate the potential of UD1022 to suppress *M. oryzae* development by inhibiting both spore germination and appressorium formation, emphasizing its promising role as a biocontrol agent.

**Fig 3.**
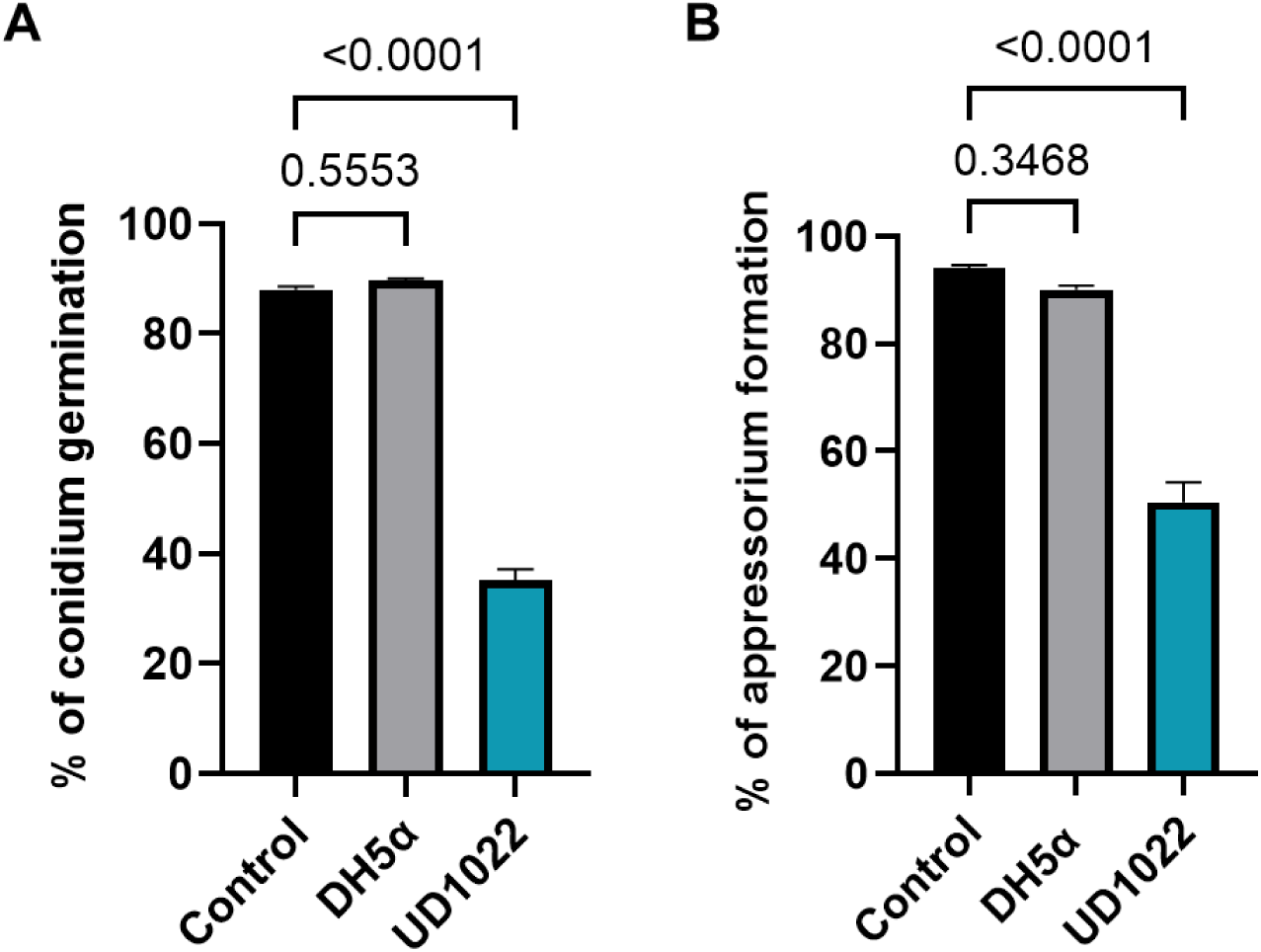
*B. subtilis* UD1022 inhibits both *M. oryzae* conidial germination and appressorium formation. **(A, B)** Effect of *B. subtilis* UD1022 on *M. oryzae* conidial germination and appressorium formation through direct bacterial treatment. Fungal spores (1 x10^5^ spores/mL) were mixed with bacterial suspensions (∼OD₆₀₀ 0.2, 1.6 × 10⁸ CFU/mL) and incubated in a 20 µL drop on an appressorium-inductive surface (hydrophobic coverslip). Control treatments included sterile water-control and *E. coli* DH5α. Conidial germination was assessed at 3 hpi, and appressorium formation was evaluated at 20 hpi. Bar graphs represent the percentage of *M. oryzae* conidial germination (left) and appressorium formation (right) under control conditions (black, gray) and in the presence of *B. subtilis* UD1022 (blue). Error bars indicate standard deviation. Statistical significance was determined using one-way ANOVA (p < 0.05).

### UD1022 primes rice root resistance against *M. oryzae*

To assess the efficacy of UD1022 in controlling *M. oryzae* infection, we conducted a bacterial inoculation followed by a pathogenicity assay to measure disease severity on rice plants. This assay was designed to determine whether UD1022 could reduce lesion development caused by *M. oryzae*. Roots of rice plants, including the moderately susceptible cultivar CO39 and the highly susceptible cultivar YT16, were treated with UD1022 suspensions and, after 24 hours, challenged with *M. oryzae* spores on the leaves (**Fig. 4**). Disease symptoms were quantified by measuring the percentage of disease-covered leaf area using ImageJ software. In our study, UD1022-treated plants showed a disease-covered area of 10%, significantly lower than the untreated controls inoculated with *M. oryzae* Guy11 or plants treated with *E. coli* DH5α, both of which exhibited an average of 60% disease-covered area in both CO39 and YT16 cultivars (**Fig. 4A-D**). Plants inoculated with UD1022 displayed significantly fewer leaf lesions compared to the untreated *M. oryzae* Guy11 control (**Fig. 4A,B**). While the control leaves showed extensive necrotic lesions, characterized by large, irregular spots covering a significant portion of the leaf surface, UD1022-treated leaves had a markedly reduced lesion number. These findings suggest that root colonization by UD1022 activates plant-driven mechanisms that prime rice for foliar defense against *M. oryzae*, leading to an enhanced immune response and reduced infection in the aerial parts of the plant.

**Fig 4.**
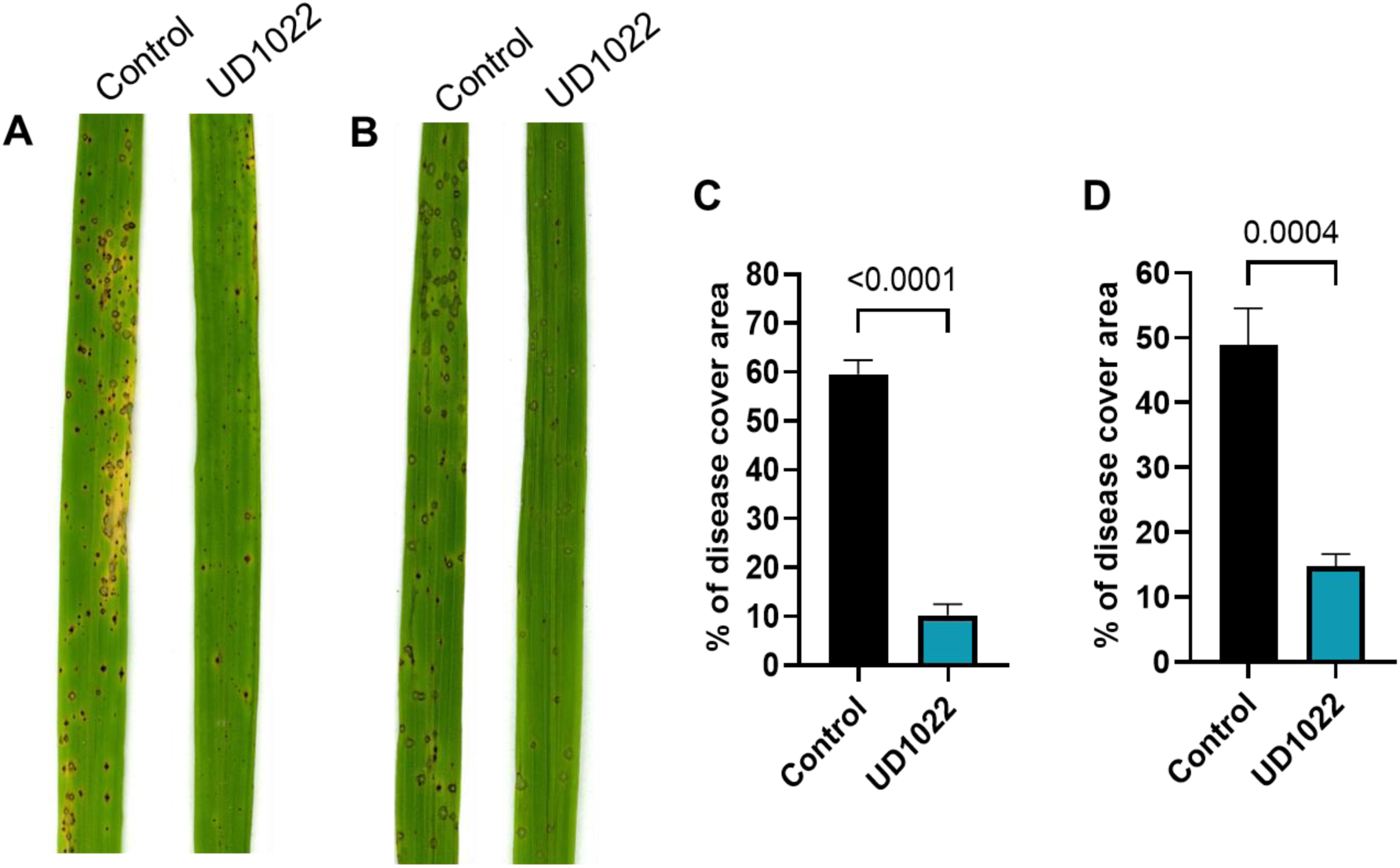
*B. subtilis* UD1022 primes rice root resistance against *M. oryzae.* **(A, B)** Representative rice leaf segments showing disease symptoms in mock-treated and *B. subtilis* UD1022-treated plants. Rice plants were root-primed with bacterial suspension (∼OD₆₀₀ 0.1, 0.8 × 10⁸ CFU/mL). After 24 hpi, CO39 (A) and YT16 (B) were spray-inoculated with an *M. oryzae* spore suspension (1 × 10⁵ spores/mL). **(C, D)** Quantification of disease severity, represented as the percentage of leaf area covered by lesions in CO39 (C) and YT16 (D) rice cultivars. Bar graphs show the percentage of *M. oryzae* appressorium formation under control (mock-treated, black) and *B. subtilis* UD1022-treated (blue) conditions. Error bars represent standard deviation. Statistical significance was determined using Student’s *t*-test (*p* < 0.05).

### Defense gene activation in rice by *B. subtilis* UD1022

To further investigate the role of UD1022 in enhancing rice defense against *M. oryzae*, we examined the expression of key defense-related genes 24 hours post-bacterial treatment (**Fig. 5; Table S1**). We focused on the salicylic acid (SA), jasmonic acid (JA), and ethylene (ET) pathways due to their central roles in plant immunity: SA is crucial for defense against biotrophic pathogens, JA mediates responses to necrotrophic pathogens and herbivory, and ET regulates stress signaling and cross-talk between pathways (27). We selected genes previously identified as being activated in rice and *Arabidopsis* during pathogen infection and involved in ISR triggered by beneficial PGPRs, using them as molecular markers to evaluate UD1022’s impact on hormonal defense signaling. Following UD1022 treatment, there was significant upregulation of SA-responsive genes (*PAD4, EDS1,* and *WRKY62*), JA-responsive genes (*WRKY30, JAR1,* and *WRKY77*), and ET-responsive genes (*EIN2, EIL1,* and *ERF1*), indicating activation of multiple defense pathways (**Fig. 5A-C**). Notably, these changes in gene expression were observed in the absence of pathogen challenge, indicating that UD1022 alone is sufficient to activate systemic defense responses in rice. The moderate upregulation of pathogenesis-related genes (*e.g., PR10b, PAD4*) suggests that UD1022 induces a primed immune state, where the plant’s defense mechanisms are preconditioned for a faster and stronger response upon pathogen attack (**Fig. 5D**). Furthermore, the upregulation of WRKY transcription factors in UD1022-treated plants suggests that UD1022 modulates transcriptional reprogramming of defense networks, reinforcing its role in enhancing systemic resistance.

**Fig 5.**
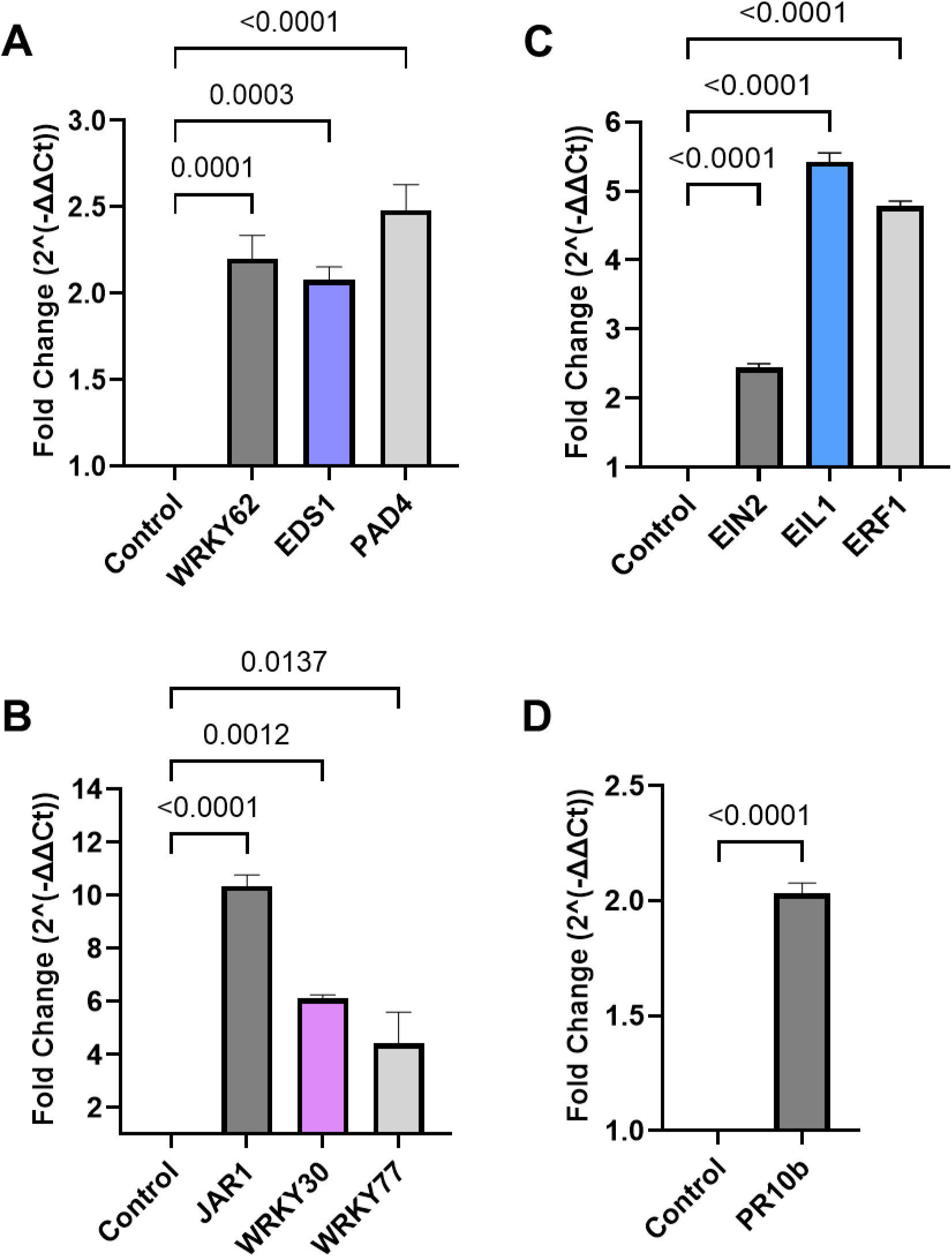
Expression of defense-related genes in rice plants treated with *B. subtilis* UD1022. Roots of aseptically grown rice plants were treated with *B. subtilis* UD1022 and water-control. Leaf samples were collected 24 hours post-treatment, and the expression of genes involved in (**A**) salicylic acid (SA), (**B**) jasmonic acid (JA), (**C**) ethylene (ET) signaling pathways, or (**D**) general defense response was analyzed via qPCR. Samples were normalized against housekeeping genes and then to water-control. Error bars represent standard deviation. Statistical significance was determined using Student-t’s test and one-way ANOVA and Student’s *t*-test (*p* < 0.05).

Taken together, rice plants root-inoculated with UD1022 exhibited a significant upregulation of defense-related genes compared to untreated plants, indicating the activation of key immune networks essential for disease resistance. This strong induction further supports the involvement of multiple defense pathways, illustrating UD1022’s ability to enhance systemic immunity. These findings suggest that UD1022 primes the plant’s immune system by activating SA, JA, and ET signaling pathways, thereby strengthening the plant’s capacity to combat *M. oryzae*. The multifaceted defense response observed in UD1022-treated plants underscores its potential as a biocontrol agent for sustainable rice disease management.

## DISCUSSION

Rice blast disease, caused by the devastating pathogen *M. oryzae*, poses a significant threat to global food security by causing substantial yield losses in rice cultivation. Sustainable management strategies are needed to reduce reliance on synthetic fungicides, which present environmental risks and drive pathogen resistance. This study emphasizes the potential of *B. subtilis* UD1022 as a biocontrol agent capable of suppressing rice blast symptoms through both antagonism activity and activation of plant defense mechanisms. To our knowledge, this is the first report demonstrating UD1022’s ability to control rice blast through these dual mechanisms. In addition to inhibiting *M. oryzae* vegetative growth, UD1022 also reduces its pathogenicity by suppressing appressorium formation.

Our dual culture assays demonstrated that UD1022 significantly inhibited *M. oryzae* vegetative growth, reducing fungal colony diameter. UD1022 was previously identified as a PGPR with demonstrated antagonistic activity against oomycetes and several fungal pathogens affecting turfgrass, alfalfa, and *Arabidopsis* plants (23, 25, 26). This Gram-positive bacterium readily forms biofilm and produces a diverse range of secondary metabolites, both of which contribute to its biocontrol potential. Efficient biofilm formation, coupled with surfactin production, appears to be crucial for direct antagonistic interactions. Previous studies link UD1022’s antifungal activity to nonribosomal peptide (NRP) production and biofilm formation. Surfactin, synthesized by srfAC, partially contributed to *Ascochyta medicaginicola* inhibition, while additional sfp-dependent metabolites likely targeted *Phytophthora medicaginis* (26). However, surfactin alone failed to inhibit *P. medicaginis*, suggesting the involvement of other antimicrobial compounds. Furthermore, biofilm-deficient mutants exhibited reduced antagonism, underscoring the importance of surface colonization in pathogen suppression. The global regulator Spo0A was found to be essential for full antagonistic activity, orchestrating NRP synthesis, biofilm formation, and motility. These mechanisms could similarly contribute to UD1022’s inhibition of *M. oryzae* in its vegetative form, though further research is needed. Bacterial volatile compounds (BVCs) are low molecular weight compounds capable of diffusing over long distances and influencing plant growth and defense responses (28). Many *Bacillus* species produce BVCs with antifungal activity, though their effectiveness varies among pathogens. In this study, volatile compound detection was performed using a stacking plate assay, revealing that *B. subtilis* UD1022 emitted VOCs that inhibited *M. oryzae* growth on plates. This finding contrasts with previous observations in other plant-pathogen systems, where UD1022 inhibited fungal growth of *C. jacksonii*, the causative agent of dollar spot disease in turfgrass, with no effect from bacterial volatiles or cell-free filtrates, indicating that direct contact with live cells is required (25). These results suggest that the effect of UD1022-derived VOCs may be pathogen-specific, indicating the need for additional studies to identify the types of VOCs involved and their role in fungal inhibition. Additionally, further research is needed to determine the exact mechanism by which VOCs reduce *M. oryzae* mycelial growth in the stacking assay and whether this inhibition affects the formation or function of appressoria, a specialized infection structure. Since appressoria formation is fully mature by approximately 20 hpi, it is also important to investigate whether VOCs require a longer exposure time beyond 20 hpi to accumulate to inhibitory levels.

In this study, we found that UD1022 disrupted spore germination and appressorium formation - two critical stages in the *M. oryzae* infection cycle - on a surface conducive to appressorium development. The appressorium is a specialized structure essential for host penetration and colonization. By inhibiting these processes, UD1022 directly interferes with the pathogen’s ability to establish infection. Similar inhibitory effects have been observed with other *Bacillus* species and biocontrol agents, revealing the importance of targeting early infection stages to mitigate disease progression (29–32). Previous studies have demonstrated that *B. subtilis* disrupts cell wall integrity and inhibits appressorium formation in *M. oryzae*, effectively suppressing rice blast disease. For instance, *B. subtilis* strain G5 disrupts mycelial integrity, inhibits fungal growth, and interferes with appressorium formation, thereby reducing the pathogenicity of *M. oryzae* (29). Similarly, *Pseudomonas chlororaphis* EA105 has been shown to inhibit conidia germination and appressorium formation through direct interactions on hydrophobic surfaces (33). Further studies are needed to explore the direct effects of UD1022 on appressorium development and its underlying mechanisms.

*Bacillus* species are well-established biocontrol agents that enhance plant resistance against pathogens through multiple mechanisms. They produce antimicrobial compounds, such as lipopeptides, which directly inhibit pathogen growth. Beyond disease suppression, *Bacillus* species, including UD1022, function as PGPR by regulating plant physiology and enhancing nutrient availability. They achieve this through auxin (IAA) biosynthesis, siderophore production, and the solubilization of soil nutrients, all of which contribute to plant health and resilience (34). These traits make *Bacillus* species valuable for improving crop productivity while reducing dependence on chemical fertilizers and pesticides.

In addition to their growth-promoting properties, *Bacillus* spp. can induce systemic resistance (ISR) in plants by activating defense-related pathways. ISR is a priming mechanism that enhances the plant’s ability to respond to future pathogen attacks more rapidly and effectively. Unlike Systemic Acquired Resistance (SAR), which is triggered by pathogen attack and leads to a prolonged immune response, ISR enhances the plant’s sensitivity to defense hormones such as JA, SA, and ET, facilitating a more rapid immune response (27, 35, 36).

The activation of ISR by beneficial microbes is initiated through recognition of microbial-associated molecular patterns (MAMPs), similar to pattern-triggered immunity (PTI), but with key differences. Beneficial microbes can either modify their MAMPs to induce a weaker and transient immune response or actively suppress immune signaling through secondary metabolites, suggesting a long evolutionary history of plant-microbe symbiosis (27). In addition to hormonal priming, ISR leads to structural, epigenetic, and transcriptomic modifications, collectively strengthening plant defenses against future pathogen challenges.

In our study, treatment with UD1022 significantly reduced rice blast symptoms while up-regulating key defense-related genes, suggesting that UD1022 not only directly inhibits *M. oryzae* but also primes rice plants for enhanced resistance via ISR. Rice plants treated with UD1022 exhibited a dramatic reduction in disease severity, with a disease-covered area of only 10% compared to untreated controls. Treated plants demonstrated a reduction in the number of necrotic lesions in both rice cultivars tested. This reduction underscores UD1022’s potential as an effective biocontrol agent for rice blast.

UD1022 treatments applied to the root system effectively induced ISR, priming plant responses in the phyllosphere and enhancing resistance to pathogen invasion. Gene expression analysis revealed significant upregulation of SA-responsive genes (*WRKY62, EDS1, and PAD4*), JA-responsive genes (*JAR1, WRKY30, and WRKY77*), and ET-responsive genes (*EIL1, ERF1, and EIN2*). While PR10b has been previously identified as an SA-responsive gene, it also responds to JA signaling, indicating crosstalk between these pathways (37, 38). These findings align with previous studies demonstrating UD1022’s capacity to trigger plant defense mechanisms. For instance, previous studies have shown that UD1022 (formerly known as *B. subtilis* FB17) limits the entry of *P. syringae* DC3000 into *A. thaliana* by triggering ISR, which induces stomatal closure in light-adapted plants, thereby reducing pathogen invasion (39). The moderate upregulation of Pathogenesis-related (PR) genes (*PR10b*, *PAD4*) further supports the conclusion that UD1022 induces immune priming rather than full defense activation, allowing the plant to maintain metabolic efficiency while remaining in a heightened state of immune readiness. Therefore, this suggests that ISR does not lead to constitutive PR gene expression but rather enhances their responsiveness to future pathogen challenge (40, 41).

Integrating multiple biocontrol agents and plant growth promoters into synthetic microbial communities offers an environmentally sustainable alternative to conventional fertilizers and pesticides. However, designing a robust community capable of withstanding diverse environmental challenges remains a significant hurdle (42). A major challenge lies in translating laboratory-based findings to field conditions, where climatic variability, nutrient fluctuations, and increased microbial diversity influence biocontrol efficacy. These factors not only affect the interaction between the BCA and its target pathogen but also shape its relationship with the host plant and native microbiome. For instance, *Bacillus* species have been shown to either enhance or reduce soil microbial diversity post-inoculation, suggesting potential competitive interactions that may impact beneficial microbes (43). Therefore, it is essential to evaluate both the pathogen-targeting ability of BCAs and their broader ecological impact on the host microbiome.

This study establishes *B. subtilis* UD1022 as a promising biocontrol agent for managing rice blast disease. Its dual mechanisms of direct antagonism and ISR activation provide a comprehensive approach to disease suppression, reducing reliance on chemical inputs and supporting sustainable agriculture. To our knowledge, this is the first report of UD1022 reducing rice blast symptoms. Further research should validate UD1022’s efficacy in field conditions, explore its interactions with native soil microbiota, and optimize its formulation for sustainable, large-scale application in rice blast management.

## CONCLUSION

In this study, we demonstrate that *B. subtilis* UD1022 effectively suppresses *M. oryzae* growth and infection through multiple mechanisms, including direct antagonism, VOC-mediated inhibition, and the induction of systemic resistance in rice plants. Dual culture and stacking plate assays confirmed UD1022’s ability to inhibit fungal growth, while in planta experiments showed a significant reduction in disease severity and lesion formation in treated plants. Gene expression analysis revealed the upregulation of key defense-related genes in the SA, JA, and ET signaling pathways, indicating a primed immune response. Collectively, these findings demonstrate UD1022 as a potent biocontrol agent with the potential to be integrated into sustainable rice blast management strategies.

## MATERIAL AND METHODS

### Fungal and bacterial cultures

The fungal strain *M. oryzae* Guy11, a model for plant-pathogen interaction studies, was cultured on Complete Media (CM) agar (50 mL 20× Nitrate Salts, 1 mL Trace Elements, 10 g D-glucose, 2 g Peptone, 1 g Yeast Extract, 1 g Casamino Acids, 1 mL Vitamin Solution, and distilled water to 1 L, pH adjusted to 6.5 with NaOH and solidified with 15 g agar before autoclaving at 121°C for 20 minutes). The mycelial plugs were grown in CM and incubated at 24°C for a 12-hour light/dark cycle.

The bacterial strain *Bacillus subtilis* UD1022, a plant growth-promoting rhizobacterium, was provided by Dr. Harsh Bais (University of Delaware). Glycerol stocks were stored at −80°C and revived on LB agar (10 g/L Tryptone, 5 g/L Yeast Extract, 5 g/L NaCl, and 18 g/L Agar, pH ∼7.0). Plates were incubated at 30°C overnight, and single colonies were grown in liquid LB medium under shaking at 200 rpm.

### Dual culture assay to assess direct antagonism

Overnight bacterial cultures were grown in liquid CM at 28°C with shaking at 220 rpm and normalized to an OD₆₀₀ of 0.5 using sterile water. Fungal cores (5 mm), collected from the edges of 7-day-old fungal plates with flame-sterilized tools, were transferred to CM plates and placed 1.5 cm from the center. A 5 µL aliquot of normalized culture was added 3 cm away on the opposite side, with sterile water used as a negative control. Plates were sealed with parafilm and incubated at 25°C in the dark for 5 days. Fungal growth diameters were measured, averaged, and used to calculate the percentage of inhibition using the formula: % of inhibition = [(D1 - D2)/(D1)] × 100. D1= average control diameter, D2= average treatment diameter.

### Volatile-mediated assay

The volatile-mediated assay uses a stacking plate approach to evaluate the inhibitory effects of volatile organic compounds produced by bacterial isolates. CM media and bacterial cultures were prepared as described in the dual culture assay. CM plates were inoculated with 5 µL of normalized bacterial culture, spread with sterile glass beads, and incubated overnight at 28°C in the dark. Sterile water served as a negative control. Fungal cores, collected from the edges of 7-day-old fungal cultures using sterilized tools, were transferred to the center of fresh CM plates. The lid of the bacterial plate was removed, and the bacterial plate was inverted over the fungal plate to create a sealed chamber secured with parafilm. The plates were incubated for 5 days at 25°C in the dark. Fungal growth diameters were measured across the widest points, averaged, and used to calculate the percentage of inhibition as in the dual culture experiment.

### Plant material and growth conditions

*Oryza sativa* susceptible cultivars, CO39 and YT16, were gifts from Dr. Richard Wilson (University of Nebraska) and Dr. Martin Egan (University of Arkansas), respectively. Dehusked rice seeds were surface sterilized with 70% ethanol (1 min) and 5% bleach (1 hour), rinsed with sterile water, dried, and planted in the Magenta boxes. Seeds were pre-germinated on Half-strength Murashige and Skoog (MS) media (2.25 g/L MS medium, 30 g/L sucrose, and 0.5 g/L MES. pH 5.8 using KOH, and 3 g/L gelrite). Magenta boxes were filled with 100 mL of sterilized media. Plants were grown for 10 days in a growth chamber at 25°C with a 12-hour light/dark cycle. After 10 days, plants were removed from MS media and transplanted into surface-sterilized 4-inch pots filled with autoclaved soil and transferred to the greenhouse.

### Fungal spore germination and appressorium formation

Spores from *M. oryzae* Guy11 were harvested from 7–10-day-old CM agar cultures, suspended in water, and filtered through two layers of sterile Miracloth. Sterilized plastic coverslips served as hydrophobic surfaces for the conidia. A spore suspension (1×10⁵ spores/mL) was prepared, and bacterial suspensions (OD₆₀₀ = 0.02, ∼1×10⁷ cells/mL) were used for treatments. Fungal spores were mixed with bacterial suspensions at a 1:1 ratio, and 20 µL were placed in the coverslip. Control treatments included *E. coli* DH5α and untreated spores. Treated spores (20 µL) were placed on coverslips, which were incubated in a Pyrex dish containing a wet filter to maintain humidity and incubated at 25°C. Germination percentages were determined after 3 hours by counting germinated spores relative to the total spore count. Appressorium formation was assessed at 20 hours by counting germinated conidia that produced appressoria. Observations were made using a light microscope, and data from three biological replicates were analyzed for statistical significance.

### Infection assays

Bacterial cultures were prepared by growing samples in LB broth at 30°C and 220 rpm overnight. Cultures were normalized to OD₆₀₀ 0.1, washed twice with sterile water, and resuspended in 10 mL of sterile water. Ten-day-old plants were inoculated by submerging roots in bacterial suspensions for 20 minutes before being transplanted into sterile soil pots. At three weeks, plants were reinoculated with 10 mL of bacterial culture added to the soil surrounding each plant.

After 24 hours, plants were sprayed with an *M. oryzae* spore suspension (1 × 10⁵ spores/ml in 0.2% gelatin). To maintain high humidity, plants were covered with plastic bags for 48 hours and incubated in a growth chamber at 25°C with a 12-hour light/dark cycle. Bags were partially opened after two days to reduce humidity. After five days, infected leaves were collected for imaging. Disease symptoms were quantified by measuring the percentage of disease-covered areas using ImageJ software.

### Gene expression analysis

Following inoculation with bacterial isolates, leaves were collected for RNA extraction using a plant RNA isolation kit (RNeasy Plant Mini Kit, Qiagen), following the manufacturer’s protocol. The quality and concentration of the RNA were assessed using a spectrophotometer. First-strand complementary DNA (cDNA) was synthesized from 1 µg of total RNA using a reverse transcription kit with oligo-dT primers (QuantaBio).

Quantitative PCR (qPCR) was performed using a SYBR Green-based detection system in a thermal cycler (BioRad). Reactions were conducted in triplicate for each sample, with a total volume of 20 µL per reaction, containing cDNA, primers specific to the target gene, and SYBR Green master mix (QuantaBio). Amplification conditions included an initial denaturation step, followed by 40 cycles of denaturation, annealing, and extension at optimized temperatures. Expression levels of the target genes were normalized against the housekeeping gene *GAPDH*. The comparative CT method (*ΔΔCT*) was used to calculate relative gene expression levels compared to an uninoculated control. Data from three biological replicates were included for statistical analysis. Graphs and statistical analyses were performed using GraphPad Prism software. Results were visualized as fold changes in gene expression, with statistically significant differences determined using appropriate tests (*p* < 0.05). Error bars represent the standard deviation of biological replicates.

### Statistical analysis

Statistical analysis was performed using Student’s *t*-test for comparisons between two conditions and one-way ANOVA for comparisons across multiple conditions. Except for the *in planta* root inoculations, all assays included a minimum of three biological replicates. Results were considered statistically significant when the *p*-value was below 0.05. Graphical representations of significant data are annotated as follows: * (*p* ≤ 0.05), ** (*p* ≤ 0.01), *** (*p* ≤ 0.001), and **** (*p* ≤ 0.0001).

## ACKNOWLEDGEMENTS

We apologize to our colleagues whose work was not included due to space limitations or the focus of the article; we greatly appreciate their important contributions to the field. This work was supported by the University of Florida’s Research Opportunity Seed Fund Award. Special thanks to Fernandez’s laboratory for reviewing the manuscript and providing constructive feedback to improve it.

## AUTHOR CONTRIBUTIONS

Conceptualization, J.F.; Writing, J.F. and T.J.; Experiments: T.J., L.K., N.G., and J.F. Supervision, J.F. All authors contributed to the reviewing and editing of the manuscript. All authors have read and agreed to the published version of the manuscript.

## CONFLICTS OF INTEREST

The authors declare no conflict of interest.

## Supplemental Material

**Figure S1.**
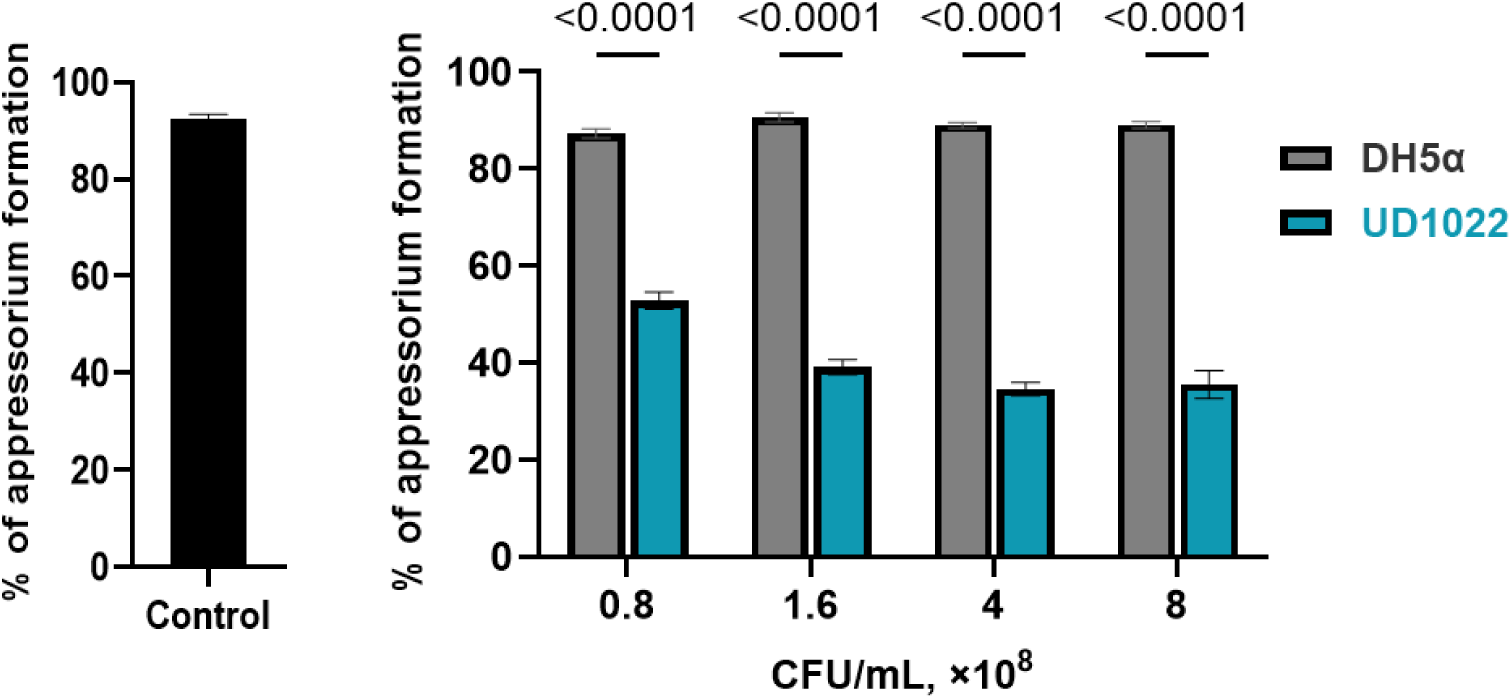
UD1022 inhibits *M. oryzae* appressorium formation at different concentrations. Effect of *B. subtilis* UD1022 on *M. oryzae* appressorium formation through different bacterial concentrations. Fungal spores (1 x10^5 spores/mL) were mixed with different bacterial suspensions and incubated in a 20 µL drop on an appressorium-inductive surface (hydrophobic coverslip). Appressorium formation was evaluated at 20 hpi. Bar graphs represent the percentage of *M. oryzae* appressorium formation under control conditions (black) and in the presence of UD1022 (blue) and *E. coli* DH5α. Error bars indicate standard deviation. Statistical significance was determined using one-way ANOVA (p < 0.05).

**Table S1.**
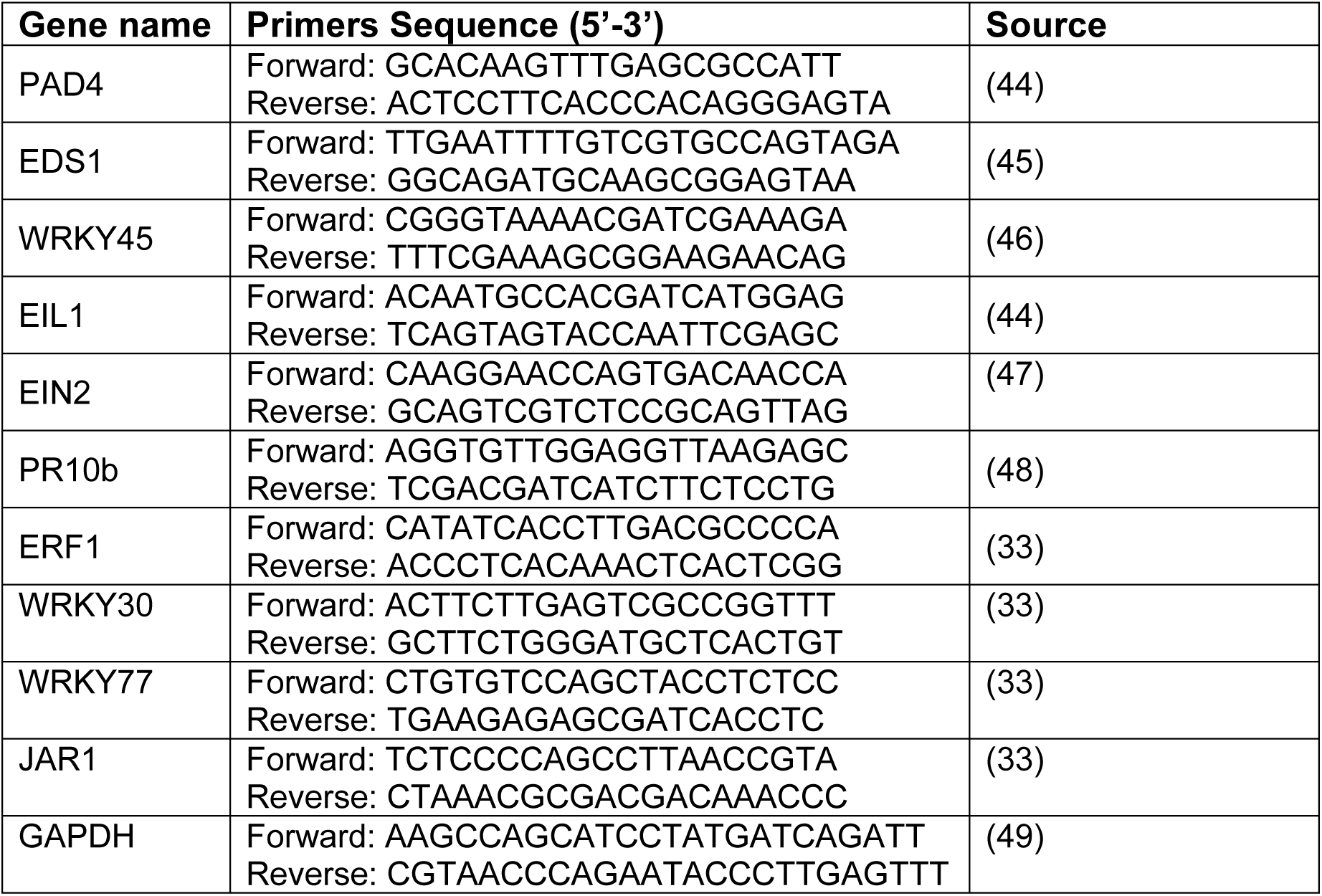
List of PCR primers used for qRT-PCR.

